# Novel DNA methylation sites of glucose and insulin homeostasis: an integrative cross-omics analysis

**DOI:** 10.1101/432070

**Authors:** Jun Liu, Elena Carnero-Montoro, Jenny van Dongen, Samantha Lent, Ivana Nedeljkovic, Symen Ligthart, Pei-Chien Tsai, Tiphaine C. Martin, Pooja R. Mandaviya, Rick Jansen, Marjolein J. Peters, Liesbeth Duijts, Vincent W.V. Jaddoe, Henning Tiemeier, Janine F. Felix, Audrey Y Chu, Daniel Levy, Shih-Jen Hwang, Jan Bressler, Rahul Gondalia, Elias L. Salfati, Christian Herder, Bertha A. Hidalgo, Toshiko Tanaka, Ann Zenobia Moore, Rozenn N. Lemaitre, Min A. Jhun, Jennifer A. Smith, Nona Sotoodehnia, Stefania Bandinelli, Luigi Ferrucci, Donna K. Arnett, Harald Grallert, Themistocles L. Assimes, Lifang Hou, Andrea Baccarelli, Eric A Whitsel, Ko Willems van Dijk, Najaf Amin, André G. Uitterlinden, Eric J.G. Sijbrands, Oscar H. Franco, Abbas Dehghan, Tim D. Spector, Josée Dupuis, Marie-France Hivert, Jerome I. Rotter, James B. Meigs, James S. Pankow, Joyce B.J. van Meurs, Aaron Isaacs, Dorret I. Boomsma, Jordana T. Bell, Ayşe Demirkan, Cornelia M. van Duijn

## Abstract

Despite existing reports on differential DNA methylation in type 2 diabetes (T2D) and obesity, our understanding of the functional relevance of the phenomenon remains limited. Because obesity is the main risk factor for T2D and a driver of methylation from previous study, we aimed to explore the effect of DNA methylation in the early phases of T2D pathology while accounting for body mass index (BMI). We performed a blood-based epigenome-wide association study (EWAS) of fasting glucose and insulin among 4,808 non-diabetic European individuals and replicated the findings in an independent sample consisting of 11,750 non-diabetic subjects. We integrated blood-based *in silico* cross-omics databases comprising genomics, epigenomics and transcriptomics collected by BIOS project of the Biobanking and BioMolecular resources Research Infrastructure of the Netherlands (BBMRI-NL), the Meta-Analyses of Glucose and Insulin-related traits Consortium (MAGIC), the DIAbetes Genetics Replication And Meta-analysis (DIAGRAM) consortium, and the tissue-specific Genotype-Tissue Expression (GTEx) project. We identified and replicated nine novel differentially methylated sites in whole blood (P-value < 1.27 × 10^−7^): sites in *LETM1, RBM20, IRS2, MAN2A2* genes and 1q25.3 region were associated with fasting insulin; sites in *FCRL6*, *SLAMF1, APOBEC3H* genes and 15q26.1 region were associated with fasting glucose. The association between *SLAMF1*, *APOBEC3H* and 15q26.1 methylation sites and glucose emerged only when accounted for BMI. Follow-up *in silico* cross-omics analyses indicate that the *cis*-acting meQTLs near *SLAMF1* and *SLAMF1* expression are involved in glucose level regulation. Moreover, our data suggest that differential methylation in *FCRL6* may affect glucose level and the risk of T2D by regulating *FCLR6* expression in the liver. In conclusion, the present study provided nine new DNA methylation sites associated with glycemia homeostasis and also provided new insights of glycemia related loci into the genetics, epigenetics and transcriptomics pathways based on the integration of cross-omics data *in silico*.

## Background

Type 2 diabetes (T2D) is a common metabolic disease, characterized by disturbances in glucose and insulin metabolism, that are in part genetically driven^1-10^ with the heritability ranging from 20% to 80%^11^. DNA methylation has been associated with T2D as well as with fasting glucose and insulin^12^. Methylation-based risk scores of T2D predicted incident T2D cases that go beyond traditional risk factors such as obesity and waist-hip ratio^13^. Further, obesity, which is the most important determinant of insulin resistance and glucose levels in the population,^14,15^ has also been associated with differential DNA methylation^13^. This raises the possibility that differential methylation associated with glucose and insulin levels could be counfounded by obesity. DNA methylation, mainly depending on the region, results in gene silencing and thus regulates gene expression and subsequent cellular functions^16^. It is very well possible that the epigenetic modifications occur in early phases of the pathology of T2D, requiring research focusing on the early process of the disease, e.g. in subjects free of diabetes.

We aimed to determine the association of DNA methylation with fasting glucose and insulin accounting for the effect of obesity in the non-diabetic subjects and to evaluate the impact of DNA methylation on cross-omics level. We followed the hypothesis that genetic variants drive DNA methylation which subsequently regulates gene expression and then glycemic traits, changes of which mark the early phases of diabetes pathology (**Figure 1a**). First, we performed a blood-based epigenome-wide association study (EWAS) meta-analysis of 4,808 diabetes-free individuals of European descent and replicated our findings among 11 cohorts summing up to 11,750 trans-ethnic non-diabetic individuals, mainly from European ancestry. Subsequently, we explored the role of genetics in determining the regulation of methylation associated with glycemic traits and the effects of the differential methylation on the human transcriptome *in silico* (**Figure 1a**).

**Figure 1.**
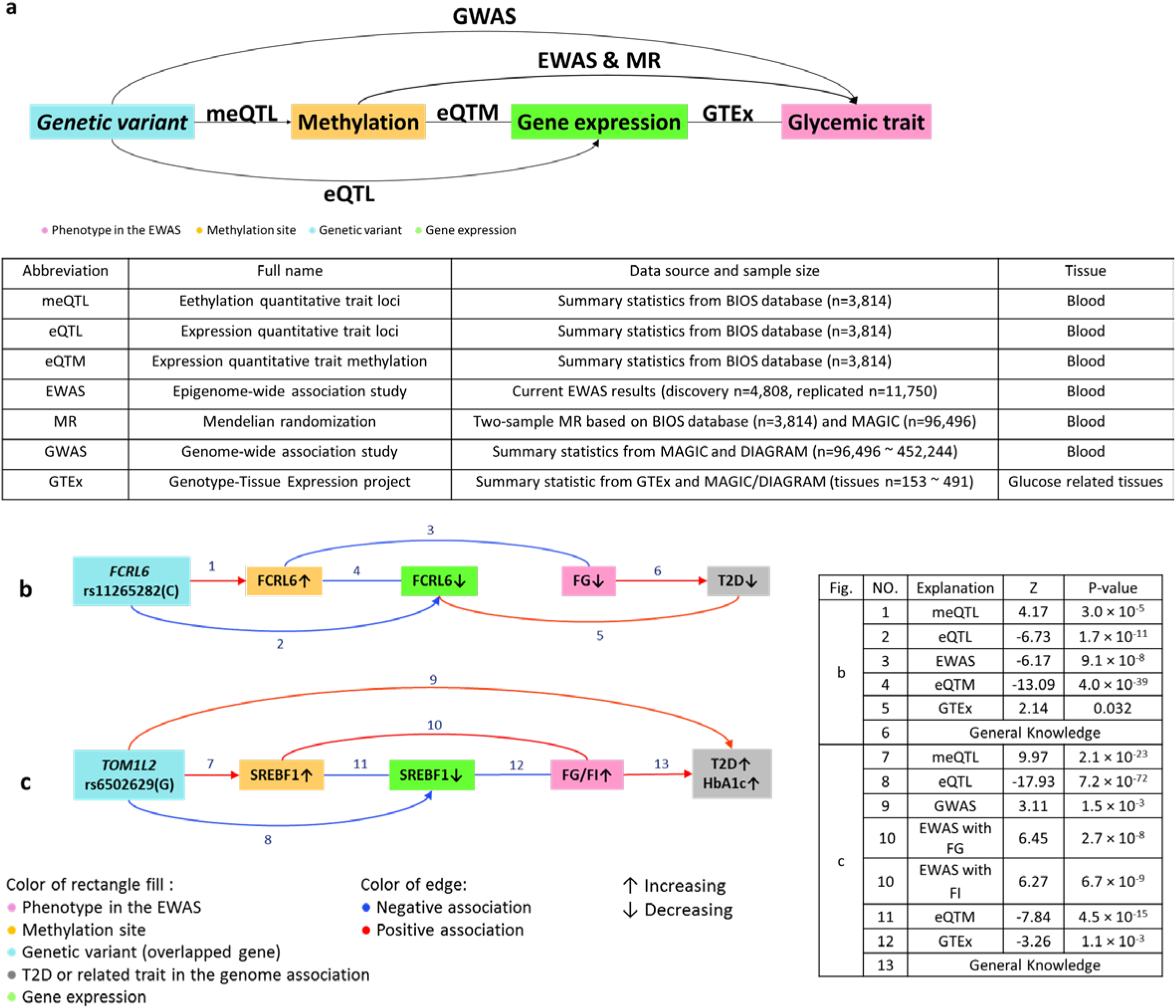
Overview of the cross-omics analysis and examples. Cascading associations cross multiple-omics-based on different data sources were integrated in the network figures. The assumption is genetic variants drive DNA methylation which subsequently regulates gene expression and then glycemic traits. FG: fasting glucose. FI: fasting insulin. T2D: type 2 diabetes

## Results

### 1. Blood-based epigenome-wide association analysis of glycemic traits

The discovery phase was based on four European cohorts (**Supplementary Table 1**). The meta-analysis revealed DNA methylation in 28 unique CpG sites associated with fasting glucose (11 CpG sites, n = 4,808) and/or insulin (20 CpG sites, n = 4,740) at epigenome-wide significance (P-value < 1.27 × 10^−7^) in either the baseline model without body mass index (BMI) adjustment or in the second model with BMI adjustment. Of these 28 CpG sites, 15 were novel (**Table 1**) while 13 were identified by earlier EWAS studies of either T2D or related traits, including glucose, insulin, hemoglobin A1c (HbA1c), homeostatic model assessment-insulin resistance (HOMA-IR) and BMI^12,13,17-25^ (**Supplementary Table 2**). Of the known CpG sites, three located in *SLC7A11, CPT1A,* and *SREBF1* associated with both glucose and insulin. The remaining ten CpG sites associated with insulin and were located in genes *ASAM, DHCR24, RNF145, KDM2B, MYO5C, TMEM49, CPT1A, two in ABCG1* and one in the 4p15.33 region.

**Table1.** Epigenome-wide association study (EWAS) results: novel differentially methylated sites associated with fasting glucose or insulin at an epigenome-wide significance level.

| Locus | CpG | Chr | Position | Regulatory feature | Trait (s) | Discovery phase (EA) |  |  |  | Replication phase (EA+AA+HS) |  |  |  |
| --- | --- | --- | --- | --- | --- | --- | --- | --- | --- | --- | --- | --- | --- |
|  |  |  |  |  |  | Model 1 |  | Model 2 |  | Model 1 |  | Model 2 |  |
|  |  |  |  |  |  | Z | P-value* | Z | P-value* | Z | P-value | Z | P-value |
| *FCRL6* | cg00936728 | 1 | 159772194 | NA | Glucose | **-6.17** | **$9.1 \times 10^{-8}$** | -5.71 | $1.9 \times 10^{-7}$ | **-3.90** | **$9.55 \times 10^{-5}$** | NP | NP |
| *SLAMF1* | cg18881723 | 1 | 160616870 | Promoter associated | Glucose | **6.11** | **$7.5 \times 10^{-8}$** | **6.94** | **$3.4 \times 10^{-10}$** | 2.67 | $7.66 \times 10^{-3}$ | **3.23** | **$1.2 \times 10^{-3}$** |
| *1q25.3* | cg13222915 | 1 | 184598594 | | Insulin | **-6.50** | **$2.6 \times 10^{-9}$** | -4.61 | $4.1 \times 10^{-6}$ | **-8.16** | **$3.33 \times 10^{-16}$** | NP | NP |
| *BRE* | cg20657709 | 2 | 28509570 | NA | Glucose | -5.46 | $2.7 \times 10^{-6}$ | **-5.88** | **$4.1 \times 10^{-8}$** | NP | NP | -2.10 | 0.036 |
| *LRPPRC* | cg01913188 | 2 | 44223249 | Promoter associated | Glucose | 5.13 | $9.4 \times 10^{-6}$ | **6.27** | **$5.7 \times 10^{-9}$** | NP | NP | 0.12 | 0.90 |
| *IRAK2* | cg14527942 | 3 | 10276383 | NA | Insulin | **6.97** | **$3.4 \times 10^{-10}$** | **6.69** | **$2.9 \times 10^{-11}$** | -0.70 | 0.48 | -0.75 | 0.45 |
| *LETM1* | cg13729116 | 4 | 1859262 | Promoter associated | Insulin | **5.95** | **$4.3 \times 10^{-8}$** | 4.56 | $4.5 \times 10^{-6}$ | **4.96** | **$6.98 \times 10^{-7}$** | NP | NP |
| *RBM20* | cg15880704 | 10 | 112546110 | NA | Insulin | **6.41** | **$3.8 \times 10^{-9}$** | 1.38<br>4.06 | $6.7 \times 10^{-5}$ | **6.83** | **$8.62 \times 10^{-12}$** | NP | NP |
| *IRS2* | cg25924746 | 13 | 110432935 | Promoter associated | Insulin | **6.59** | **$3.0 \times 10^{-9}$** | 4.55 | $4.9 \times 10^{-6}$ | **6.65** | **$3.01 \times 10^{-11}$** | NP | NP |
| *SPTB* | cg07119168 | 14 | 65225253 | NA | Glucose | -5.86 | $4.4 \times 10^{-7}$ | **-5.82** | **$4.9 \times 10^{-8}$** | NP | NP | -1.81 | 0.070 |
| *15q26.1* | cg18247172 | 15 | 91370233 | NA | Glucose | -5.25 | $4.9 \times 10^{-6}$ | **-5.90** | **$2.8 \times 10^{-8}$** | NP | NP | **-3.47** | **$5.1 \times 10^{-4}$** |
| *MAN2A2* | cg20507228 | 15 | 91460071 | Promoter associated (Cell type specific) | Insulin | **5.90** | **$5.5 \times 10^{-8}$** | 4.83 | $9.0 \times 10^{-7}$ | **7.93** | **$2.28 \times 10^{-15}$** | NP | NP |
| *FAM92B* | cg06709610 | 16 | 85143924 | NA | Insulin | **6.35** | **$6.5 \times 10^{-9}$** | **7.24** | **$5.8 \times 10^{-13}$** | 0.24 | 0.81 | 0.55 | 0.59 |
| *CD300A* | cg08087047 | 17 | 72461209 | NA | Glucose | -5.19 | $5.9 \times 10^{-6}$ | **-5.80** | **$1.1 \times 10^{-7}$** | NP | NP | -1.08 | 0.28 |
| *APOBEC3H* | cg06229674 | 22 | 39492189 | NA | Glucose | -5.40 | $1.8 \times 10^{-6}$ | **-5.86** | **$4.7 \times 10^{-8}$** | NP | NP | **-4.82** | **$1.4 \times 10^{-6}$** |

Genome-wide DNA methylation sites were tested for association with fasting glucose or fasting insulin in two models. Novel epigenome-wide significant ( $P\text{-value} < 1.27 \times 10^{-7}$ ) results in the discovery phase and the replication are shown. Model 1 adjusted for age, sex, technical covariates, white blood cell, and smoking status, accounting for family structure if needed in each cohort. Model 2 adjusted for BMI additionally. The significant associations in non-reported CpG sites were promoted for replication of the same models and traits. NP: Replication was not performed in the non-significant associated model or trait. Locus: the cytogenetic location or the gene symbol of the CpGs from Illumina annotation. Regulatory feature: the regulatory feature group of the CpGs from Illumina annotation. Chr: Chromosome. \* Genomic controlled P-value. EA+AA+HA: European ancestry, African ancestry and Hispanic ancestry population. Z: effect estimate per standard error. **Bold print:** Significant results ( $P\text{-value} < 1.27 \times 10^{-7}$ in the discovery phase, $P\text{-value} < 3.3 \times 10^{-3}$ in the replication phase). NA: Not available.

The 15 novel CpGs were tested using the same models in meta-analysis of 11 independent cohorts including 11,750 non-diabetic subjects from the Cohorts for Heart and Aging Research in Genomic Epidemiology (CHARGE) consortium (**Supplementary Table 1**). As a result, nine unique CpG-trait associations were replicated: four passing the epigenome-wide significance threshold (P-value < 1.27 × 10^−7^) and five passing the Bonferroni significance threshold after correcting for 15 tests (P-value < 3.3 × 10^−3^) (**Table 1**). These included five sites associated with fasting insulin in the baseline model (*LETM1*, *RBM20*, *IRS2, MAN2A2* and 1q25.3 region), one associated with fasting glucose in the baseline model (*FCRL6)* and additional three emerged to be associated with fasting glucose in the BMI-adjusted model (*SLAMF1, APOBEC3H* and 15q26.1 region).

Because the replication cohorts also included other ethnic groups than the main European ancestry (European: n = 7,254, African: n = 3,744, and Hispanic: n = 543), we also performed meta-analysis stratified by ancestry. Seven out of nine new CpG sites (*FCRL6, LETM1, RBM20, IRS2, MAN2A2, APOBEC3H* and the 15q26.1 region) confirmed consistent directions of effect across the three ethnicities. (**Supplementary Table 3**)

### 2. Integrated *in silico* cross-omics studies

To evaluate the functional relevance of differential methylation findings, we integrated our EWAS findings with genomics, epigenomics and transcriptomics data obtained from public resources. These included blood-based *cis* and *trans* methylation quantitative trait loci (meQTLs), expression quantitative trait methylations (eQTMs), expression quantitative trait loci (eQTLs) from the European BIOS database^26^ from the Biobanking and BioMolecular resources Research Infrastructure of the Netherlands (BBMRI-NL), the genome-wide association studies (GWAS) of glycemic traits or T2D from the Meta-Analyses of Glucose and Insulin-related traits Consortium (MAGIC) and the DIAbetes Genetics Replication And Meta-analysis (DIAGRAM) consortium^4-7, 27^, and tissue-specific eQTL-phenotype associations from MetaXcan database^28,29^ based on Genotype-Tissue Expression (GTEx) project (See resources of these database in **URLs**). The hypotheses tested were outlined in **Figure 1a**: DNA methylation and gene expression are partly genetically driven and heritable; genetic variants determine in part methylation, which subsequently influences expression and further fasting glucose and/or insulin. While doing so, we centered on the 11 top independent DNA methylation sites previously identified (cg00574958 in *CPT1A* and cg06500161 in *ABCG1* were used) and 9 novel sites from our current study (total n = 20 methylation sites).

#### 2.1 Genomics of the differentially methylated sites involved in glycemic traits

Using BIOS database (blood-based)^26^, we found that 2,991 single-nucleotide polymorphisms (SNPs) in 29 unique genetic loci were associated with methylation in either *cis* or *trans* across 18 unique CpG sites among the tested 20 target methylation sites. For two CpG sites located in *SLC7A11* and *LETM1*, we did not find any significant meQTLs. Results are shown in **Figure 2** and given in detail in **Supplementary Table 4**. Seven of the 29 meQTLs, 5 *cis-acting* and 2 *trans-acting* were found significantly associated with T2D, fasting glucose or HbA1c (shown in **Figure 2** and given in detail **Supplementary Table 5**).

**Figure 2.**
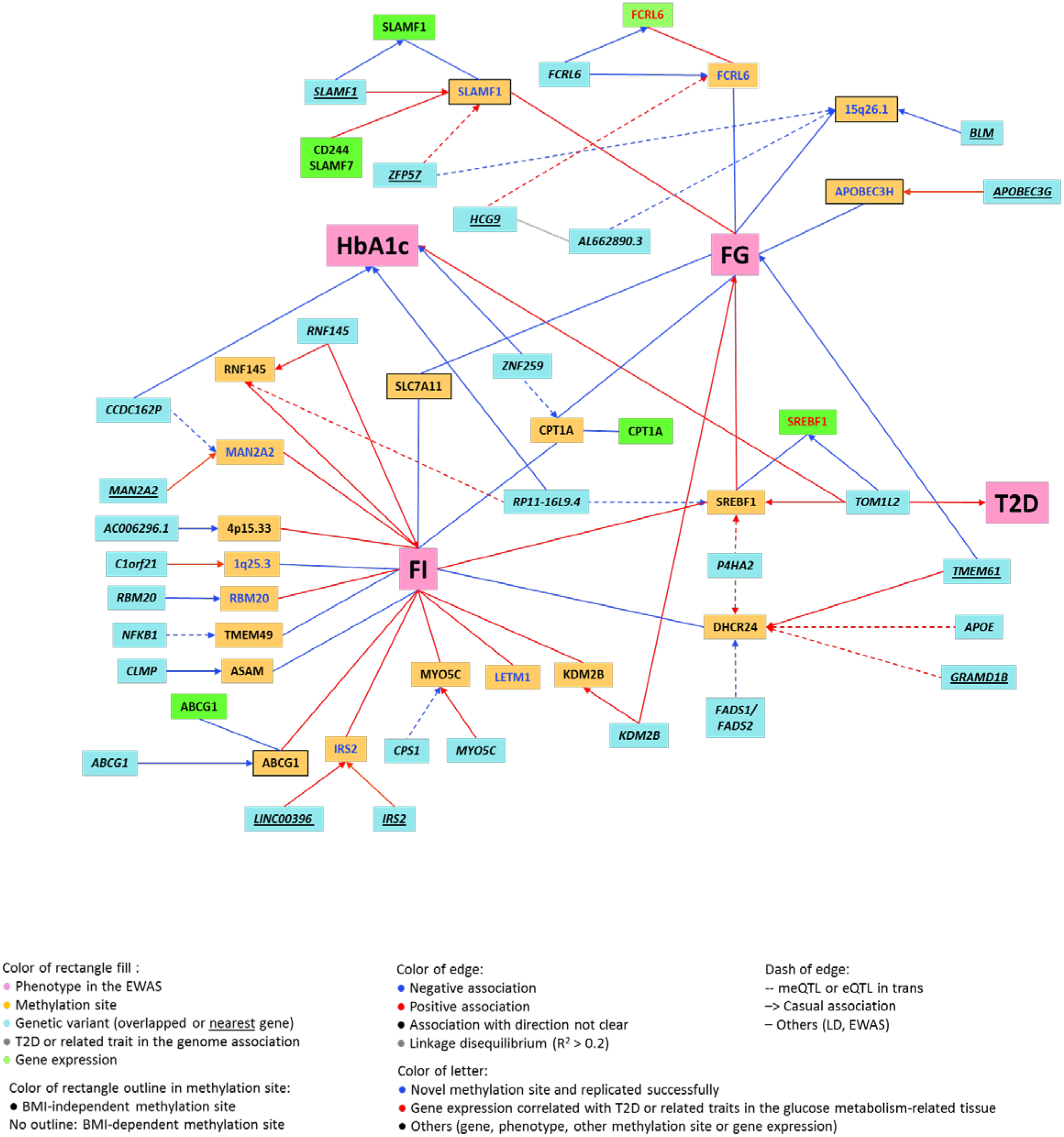
Associations between genetic variants, DNA methylation sites, gene expressions and fasting glucose, insulin and related traits based on the integration of cascading associations in Figure 1a. The effect allele is standardized across all associations. Only the significant associations which passed the specific P-value threshold in each association step were shown in the figure. FG: fasting glucose. FI: fasting insulin. T2D: type 2 diabetes.

Based on our leading hypothesis, we examined whether DNA methylation may influence fasting glucose and insulin in the circulation. To this end, we performed a two-sample-based Mendelian Randomization (MR) analyses^30^ to examine the causal effect of the differential DNA methylation sites in blood on fasting glucose or insulin using the summary GWAS results from BIOS^26^ and MAGIC databases^5^ (**Supplementary Table 6**). Up to eight independent genetic variants were included in the genetic risk score as the instrumental variable of each methylation site to check the association with the observationally associated traits, either fasting glucose or fasting insulin. Thirteen CpG sites out of the initial 20 met the present MR criteria and were testes by MR. No significant associations were detected when adjusting for multiple testing involving 13 independent tests (P-value < 3.8 × 10^−3^) except for two marginal siginificant associations between methylation site in *RBM20* with fasting insulin (P-value = 0.04) and methylation site in *SLAMF1* with fasting glucose (P-value = 0.05).

#### 2.2 Transcriptome associated with the differentially methylated sites of glycemic traits

##### 2.2.1 Association of gene expression with differentially methylated sites in blood

To understand if the methylation is also eQTM, we investigated the association between gene expression and the 20 key glycemic methylation sites from the European blood-based BIOS database^26^ (intergrated in **Figure 2**). We found that methylation in five CpG sites, including two novel sites (in *FCRL6* and *SLAMF1*) and three known sites (in *CPT1A*, *SREBF1* and *ABCG1*), was significantly negatively associated with the expression of their respective genes: *FCRL6* (P-value = 4.0 × 10^−39^), *SLAMF1 (*P-value = 4.1 × 10^−5^), *CPT1A* (P-value = 3.1 × 10^−20^), *SREBF1* (P-value = 4.5 × 10^−15^) and *ABCG1* (P-value = 2.2 × 10^−37^). The methylation site in *SLAMF1* was positively associated with expression of two other genes near *SLAMF1* (*CD244:* P-value = 2.9 × 10^−6^ and *SLAMF7:* P-value = 5.4 × 10^−9^). (**Supplementary Table 7**)

##### 2.2.2 Common genetic determinants of glycemia related to DNA methylation and gene expression in blood

We next explored if the genetic variants associated with the differential expression above were the same as the meQTLs using the eQTL data from the European blood-based BIOS database^26^ (integrated in **Figure 2** and detailed in **Supplementary Table 8**). We found three genetic determinants associated with both differential DNA methylation, including two novel methylation sites (in *FCRL6* and *SLAMF1*) and one known (in *SREBF1*), and gene expression in blood. Rs1577544 near *SLAMF1* associated with decreased methylation of the cg18881723 (Z = −5.45, P-value = 5.1 × 10^−8^) as well as *SLAMF1* expession (Z = −6.40, P-value = 1.6 × 10^−10^). Rs11265282 in *FCRL6* associated with increased methylation of cg00936728 (Z = 4.17, P-value = 3.0 × 10^−5^) but decreased *FCRL6* expession (Z = −6.73, P-value = 1.7 × 10^−11^). Rs6502629 in *TOM1L2* associated with increased methylation of cg11024682 (Z = 9.97, P-value = 2.1 × 10^−23^) but decreased *SREBF1* expession (Z = −17.93, P-value = 7.2 × 10^−72^).

##### 2.2.3 Tissue-specific differential expression associated with T2D and related traits

We then explored the tissue-specific differential expression associated with T2D and related traits by data-mining from MetaXcan database from the GTEx project^28,29^. This analyses targeted on the eQTM-related expression of seven genes as shown in 2.2.1 in six glucose-metabolism-related tissues including blood, adipose subcutaneous, adipose visceral omentum, liver, pancreas, and muscle skeletal (**Supplementary Table 9**). The effect direction consistency was checked between methylation sites, gene expression and T2D or related traits. That meant the direction of the association between methylation and T2D or related traits should be a combination of the directions of methylation with gene expression and gene expression with T2D or related traits. The expression of *SREBF1* in blood was significantly associated with decreased levels of HbA1c (Z = −3.26, P-value = 1.1 × 10^−3^) and also with decreased risk of T2D (Z = −2.40, P-value = 0.016). Meanwhile, the known cg11024682 in *SREBF1* was positively associated with fasting glucose (Z = 6.45, P-value = 2.7 × 10^−8^) and fasting insulin (Z = 6.27, P-value = 6.7 × 10^−9^) and negatively associated with expression of *SREBF1* in blood (Z = −7.84, 4.5 × 10^−15^) (shown in **Figure 1c**). Higher liver gene expression of *FCRL6*, a novel locus, was associated with increased risk of T2D (Z = 2.14, P-value = 0.032) based on MetaXcan^28,29,31^ results generated by integrating functional data in liver^32,33^ and the GWAS of T2D^9^. The novel cg00936728 in *FCRL6* was negatively associated with fasting glucose (Z = −6.17, P-value = 9.1 × 10^−8^) and expression of *FCRL6* in blood (Z = −13.09, 4.0 × 10^−39^) (shown in **Figure 1b**).

## Discussion

The current large-scale EWAS identified and replicated nine new methylation sites associated with fasting glucose or insulin, including three additionally uncovered sites (in *SLAMF1*, *APOBEC3H* and the 15q26.1 region) associated with fasting glucose only after adjustment for BMI. We further validated 13 previous reported CpG sites in 11 independent loci. Based on the cross-omics analyses, our report complements earlier studies^12,13,17-25,34^ for multiple DNA methylation sites related to the pathology early in the development of T2D through genetics and/or gene expression. We also present *in silico* evidence supporting the potential involvement of the nine new methylation sites.

The novel methylation sites annotated to genes that play roles in glucose and energy metabolism (*IRS2*), metabolism of proteins (*MAN2A2* and *EDEM3*, the nearest gene of cg13222915), RNA and splicing regulation (*RBM20*), RNA metabolism (*APOBEC3H*), small molecule transport (*LETM1*) and immune system process (*SLAMF1, FCRL6* and *SV2B*, the nearest gene of cg18247172). Some of these genes are also involved in other diseases or biomarkers, including inflammatory phenotypes (*EDEM3* with systemic lupus erythematosus^35^, *SLAMF1* with inflammatory bowel disease^36^ and *FCRL6* with C-reactive protein (CRP)^37,38^), cardiovascular phenotypes (*RBM20* with electrocardiographic traits^39^), cancer (*IRS2* with prostate cancer^40^) and schizophrenia (*MAN2A2*)^41^. Thus, observations provided insight into the pathways that might link T2D to inflammation, cardiovascular disease, cancer and schizophrenia, all disorders associated epidemiologically or clinically with T2D. This phenomenon may point at genetic pleiotropy of the genes, i.e. a gene codes the same products in various cells or have cascade-like signaling function that affects various targets.

In this paper, we used the assumption that genetic variants drive DNA methylation which subsequently regulates gene expression and then glycemic traits^42^. Two pathways (on *SREBF1* and *FCRL6*) related to genetics-epigenetics-transcriptomics-phenotype were observed in the present study (**Figure 1b** and **Figure1c**). We validated the differential methylation of *SREBF1* in insulin metabolism^43^ and extended the findings building a pathway based on the cascading cross-omics analysis in the assumption of genetics-epigenetics-transcriptomics-phenotype. We also discovered a new pathway of *FCRL6* in glucose metabolism, which still needs further research for its role to be fully understood. From the present study, with the integration of all the significant associations, the effect allele (*C*) of the genetic variant rs11265282 in *FCRL6* increases the methylation level which associates with lower expression of *FCRL6* in the blood. The decreased *FCRL6* expression in liver was also associates with decreased risk of T2D. This presumably is mediated by a decrease in fasting glucose.

The present study provides new genomic targets for further work on the pathology of T2D through large-scale EWAS and replication. However, the main findings are based on data from blood which was the only accessible tissue and may not be representative of more glucose-relevant tissues, although concordance of differential methylation between blood and adipose is high for certain pathways^44^. Our present MR analyses yielded no evidence for causality between methylation sites and fasting glucose or insulin. One limitation we faced here was the limited data to perform MR in all the association steps, e.g. the association of methylation with gene expression, the gene expression with phenotypes, some CpG sites with phenotypes, as well as the inverse causal effect of glucose or insulin on DNA methylation, thus we can not exclude entirely the influence of glycemia homeostasis on methylation levels. On the other hand, some of the MR tests performed had low explained variance of the instrumental variables, i.e. seven of the 13 performed CpGs have instrumental variables explained variance less than 5%. This might partly explain the insignificant findings in MR in the current study. Further studies are needed to include additional biologically relevant tissues and perform MR based on the tissue specific meQLTs.

In conclusion, our large-scale EWAS and replication have identified nine new differentially methylated sites associated with fasting glucose or insulin. The integrative *in silico* cross-omics analysis provided new insights of both known and new glycemia related loci into the genetics, epigenetics and transcriptomics pathway. Our study suggests that the expression of seven genes associated with either glycemia related DNA methylation is altered. Two of these seven expressed genes are also associated with T2D or related traits through the tissue-specific differentical expression association analysis: one known loci in *SREBF1* and one new loci in *FCRL6*. Further biological functional experiments are requried in more directly glucose-related tissues, e.g. pancreatic cells and liver, to unravel the mechanisms.

## Online methods

### Study population

The discovery samples consisted of 4,808 European individuals without diabetes from four non-overlapped cohorts, recruited by Rotterdam Study III-1 (RS III-1, n = 626), Rotterdam Study II-3 and Rotterdam Study III-2^45^ (called as RS-BIOS, n = 705), Netherlands Twin Register^46,47^ (NTR, n = 2,753) and UK adult Twin registry^48^ (TwinsUK, n = 724). The replication sets contained up to 11,750 individuals from 11 independent cohorts from the Cohorts for Heart and Aging Research in Genomic Epidemiology (CHARGE), including up to 6,818 individuals from European ancestry, 4,355 from African ancestry and 577 from Hispanic ancestry. We excluded individuals with known diabetes, those on anti-diabetic treatment or fasting glucose ≥ 7mmol/l. Local research ethics committees approved each study, and all participants gave informed consent to each original study. The details of the cohorts and the study design are shown in **Supplementary Note.**

### Glycemic traits and covariates

Venous blood samples were obtained after an overnight fast in all discovery and replication cohorts. Details of fasting glucose and insulin measurements are shown in **Supplementary Note**. Body mass index (BMI) was calculated as weight over height squared (kg/m^2^) based on clinical examinations. Smoking status was divided into current, former and never, based on questionnaires. White blood cell counts were quantified using standard laboratory techniques or predicted from methylation data using the standard Houseman method^49^ (see **Supplementary Note** for each cohort).

### DNA methylation quantification

The Illumina Human Methylation450 array was used in all discovery and replication cohorts to quantify genome-wide DNA methylation in blood samples. We obtained DNA methylation levels reported as β values, which represents the cellular average methylation level ranging from 0 (fully unmethylated) to 1 (fully methylated). Study-specific details regarding DNA methylation quantification, normalization and quality control procedures are provided in the **Supplementary Note** and **Supplementary Table 1**.

### Epigenome-wide association analysis and replication

All statistical analyses were performed using *R* statistical software. Insulin was natural log transformed. In the discovery analysis, we first performed epigenome-wide association studies (EWAS) in each cohort separately. Linear regression analysis was used to test the association between glucose and insulin with each methylation site in the Rotterdam Study samples. Linear mixed models were used in NTR and TwinsUK accounting the family structure. We fitted two models for each cohort: 1) the baseline model adjusting for age, sex, technical covariates (chip array number and position on the array), white blood cell counts (lymphocytes, monocytes, and granulocytes) and smoking status, and 2) a second model additionally adjusting for BMI. We removed probes that have evidence of multiple mapping or contain a genetic variant in the methylation site^50^. All cohort-specific EWAS results for each model were then meta-analysed using inverse variance-weighted fixed effects meta-analysis as implemented in the “metafor” R package^51^. In total, we meta-analysed 403,011 CpGs that passed quality control in all four discovery cohorts. The detail of the quality control for each cohort could be found in the **Supplementary Note**. The association was later corrected by the genomic control factor (λ) in each meta-EWAS^52^. We produced quantile-quantile (QQ) plots of the −log_10_(*P*) to evaluate inflation in the test statistic (**Supplementary Figure 1**). A Bonferroni correction was used to correct for multiple testing and identify epigenome-wide significant results (P < 1.27 × 10^−7^). We did not correct the number of glycemic traits and models, as they are highly correlated and not independent. The genome coordinates were provided by Illumina (GRCh37/hg19). The correlation of the CpG sites located in the same gene was further checked in the overall RS III-1 and RS-BIOS samples by Pearson’s correlation test (n = 1,544) to find the independent top CpGs. For the associations discovered in the meta-EWAS that have not been reported previously, we attempted replication in independent samples using the same traits and models as in the discovery analyses. Study-specific details of replication cohorts are provided in **Supplementary Table 1** and **Supplementary Note**. Results from each replication cohort were meta-analysed using the same methods as in the discovery analyses. Bonferroni P-value < 3.3 × 10^−3^ (0.05 corrected by 15 loci tested for associations) was considered significant.

### Genomics of the differentially methylated sites and glycemic traits

We identified the genetic determinants of the significant CpG sites known or replicated through the current EWAS using the results of the *cis* and *trans* methylation quantitative trait loci (meQTLs) from European blood-based BIOS database^26^ from the Biobanking and BioMolecular resources Research Infrastructure of the Netherlands (BBMRI-NL) which captured meQTLs, expression quantitative trait loci (eQTLs) and expression quantitative trait methylations (eQTMs) from genome-wide database of 3,841 Dutch blood samples (See resources of the database in **URLs**). All the reported single-nucleotide polymorphisms (SNPs) with P-value adjusted for false discovery rate (FDR) less than 0.05 in the database were treated as the target genetic variants in the present study. The SNPs were annotated based on the information in the previous study^26^ or the nearest protein-coding gene list from SNPnexus^53,54^ on GRCh37/hg19.

We explored the associations of these DNA methylation-related SNPs with type 2 diabetes (T2D) or related traits, i.e. fasting glucose, insulin, hemoglobin A1c (HbA1c), based on public genome-wide association study (GWAS) datasets in European ancestry^4-7, 27^. We checked the effect direction consistency of the association between the SNPs, methylation sites and T2D or related traits. That is the direction of the association between SNP and T2D or related traits should be a combination of the directions of SNP with methylation sites and methylation sites with T2D or related traits. A multiple-testing correction was performed by Bonferroni adjustment (P-value < 1.8 × 10^−3^, 0.05 corrected by the 29 genetic loci shown in **Supplementary Table 4**).

For the significant CpG sites known or replicated through EWAS, we attempted to evaluate the causality effect of CpGs on their significant traits, either fasting glucose or fasting insulin, using two-sample Mendelian Randomization (MR) approach as described in detail before by Dastani *et al*^30,55,56^ based on the summary statistic GWAS results from BIOS database and the Meta-Analyses of Glucose and Insulin-related traits Consortium (MAGIC) database^5,26^ (**Supplementary Figure 2**). Briefly, we constructed a weighted genetic risk score for individual CpG on phenotype using independent SNPs as the instrument variables of the CpG, implemented in the R-package “*gtx*”. The effect of each score on phenotype was calculated as

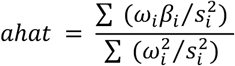
where *β_i_* is the effect of the CpG-increasing alleles on phenotype, s_*i*_ its corresponding standard error and *ω*_*i*_ the SNP effect on the respective CpG. Because the genetic variants might be close (*cis*) or far (*trans*) from the methylated site, we also performed MR test in the *cis* only SNPs if the CpG site has both *cis* and *trans* genetic markers. All SNPs were mapped to human genome build hg19. For each test (one CpG site with one trait), we extracted all the genetic markers of the CpG site in the fasting glucose or insulin GWAS from the MAGIC dataset (n = 96,496)^5^ with their effect estimate and standard error on fasting glucose or insulin. Within the overlapped SNPs, we removed SNPs in potential linkage disequilibrium (LD, pairwise R^2^ ≥ 0.05) in 1-Mbp window based on the 1000 Genome imputed genotype dataset from the general population: Rotterdam Study I (RSI, n = 6,291)^45^. We managed to exclude the genetic loci which were genome-wide significantly associated with glycemic traits, but none of the genetic loci meet this exclusion criteria. The instrumental variables that explain more than 1% of variance in exposure (DNA methylation) were taken forward for MR test. The Bonferroni P-value threshold was used to correct for the 13 CpG sites available for MR (P-value < 3.8 × 10^−3^).

### Gene expression analyses

To explore whether the differential methylation sites were associated with differential expression in blood, we explored the European blood-based BIOS database for eQTMs^26^. The significantly associated gene expression probes were searched in the eQTL data in BIOS database^26^. We also investigated if the genetic variants associated with these gene expression probes in blood were also related to the DNA methylation sites with glycemia. Finally, we tested whether the expression of the genes that harbor the eQTMs was associated with T2D and related traits in glucose metabolism-related tissues (adipose subcutaneous, adipose visceral omentum, liver, whole blood, pancreas, and muscle skeletal) using *MetaXcan*^28,29,31^ package. MetaXcan associates the expression of the genes with the phenotype by integrating functional data generated by large-scale efforts, e.g Genotype-Tissue Expression (GTEx)^32,33^ with that of the GWAS of the trait. MetaXcan is trained on transcriptome models in 44 human tissues from GTEx and is able to estimate their tissue-specific effect on phenotypes from GWAS. For this study we used the GWAS studies of T2D^9^, fasting glucose traits^5,6^, fasting insulin^6^, hemoglobin A1c (HbA1c)^57^ and homeostatic model assessment-insulin resistance (HOMA-IR)^4^. We used the nominal P-value threshold (P-value = 0.05) as we had separate assumptions for each terminal pathway between gene expressions and phenotype. Further, we checked the effect direction consistency of the association between the methylation sites and fasting glucose or insulin with the combination of the associations between the methylation sites and gene expression and between the gene expression and T2D or related traits.

### URLs

BIOS database, https://genenetwork.nl/biosqtlbrowser/; SNPnexus, http://snp-nexus.org/index.html; GWAS database of glycemic traits, https://www.magicinvestigators.org/; GWAS database of T2D, http://diagram-consortium.org/; MetaXcan, https://s3.amazonaws.com/imlab-open/Data/MetaXcan/results/metaxcan_results_database_v0.1.tar.gz. (available: 1st Jan, 2018)

---

**Supplementary Table 1.**
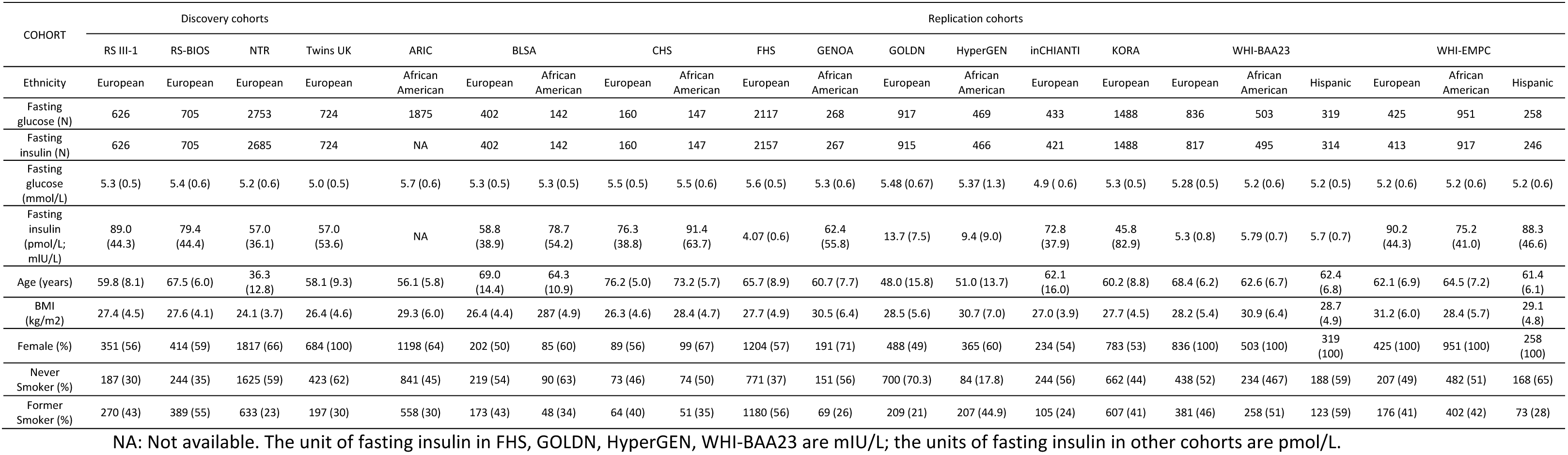
Characteristics of cohorts.

**Supplementary Table 2.**
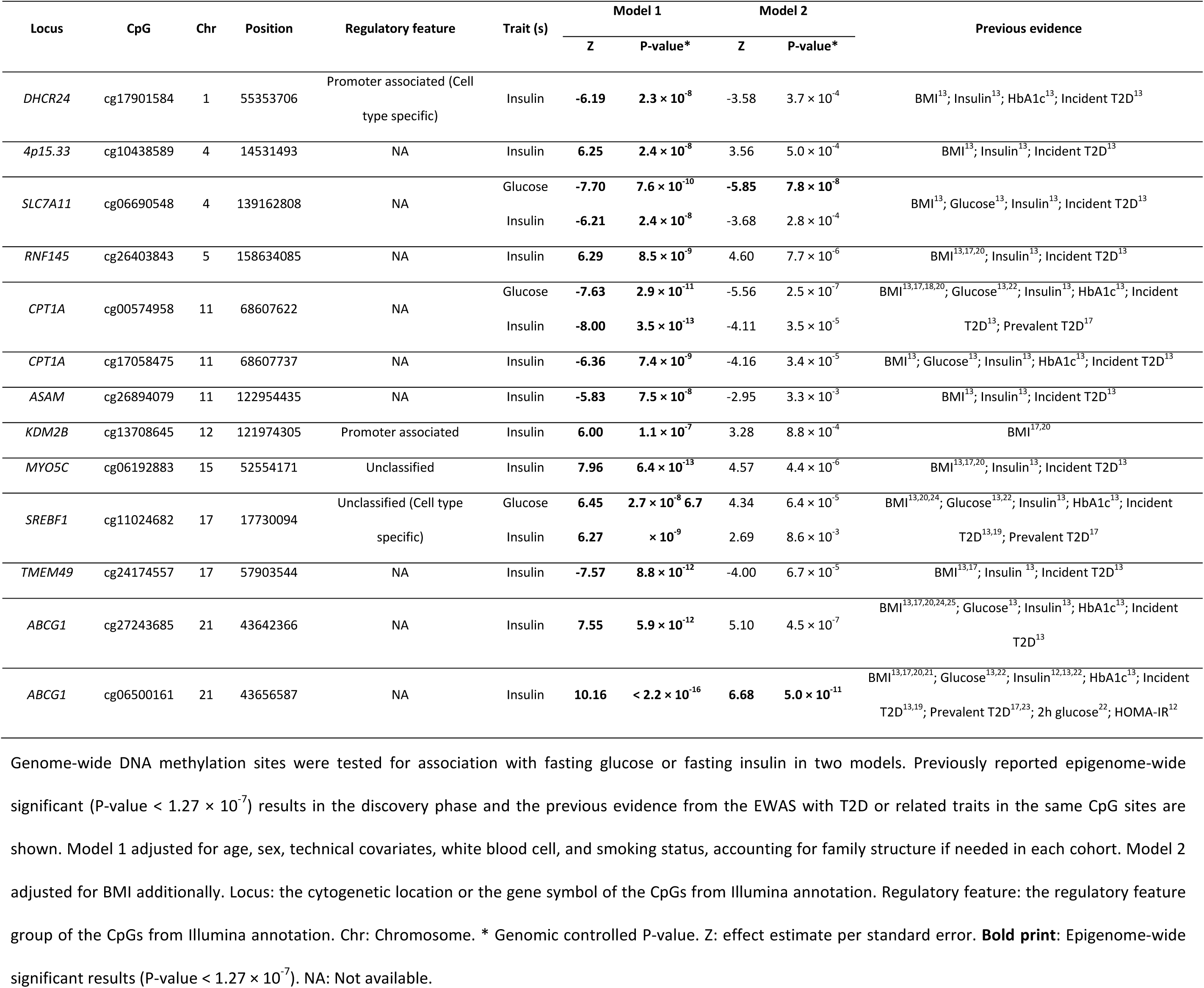
Epigenome-wide association study (EWAS) results: known differentially methylated sites associated with fasting glucose or insulin at epigenome-wide significance level in the discovery phase.

**Supplementary Table 3.**
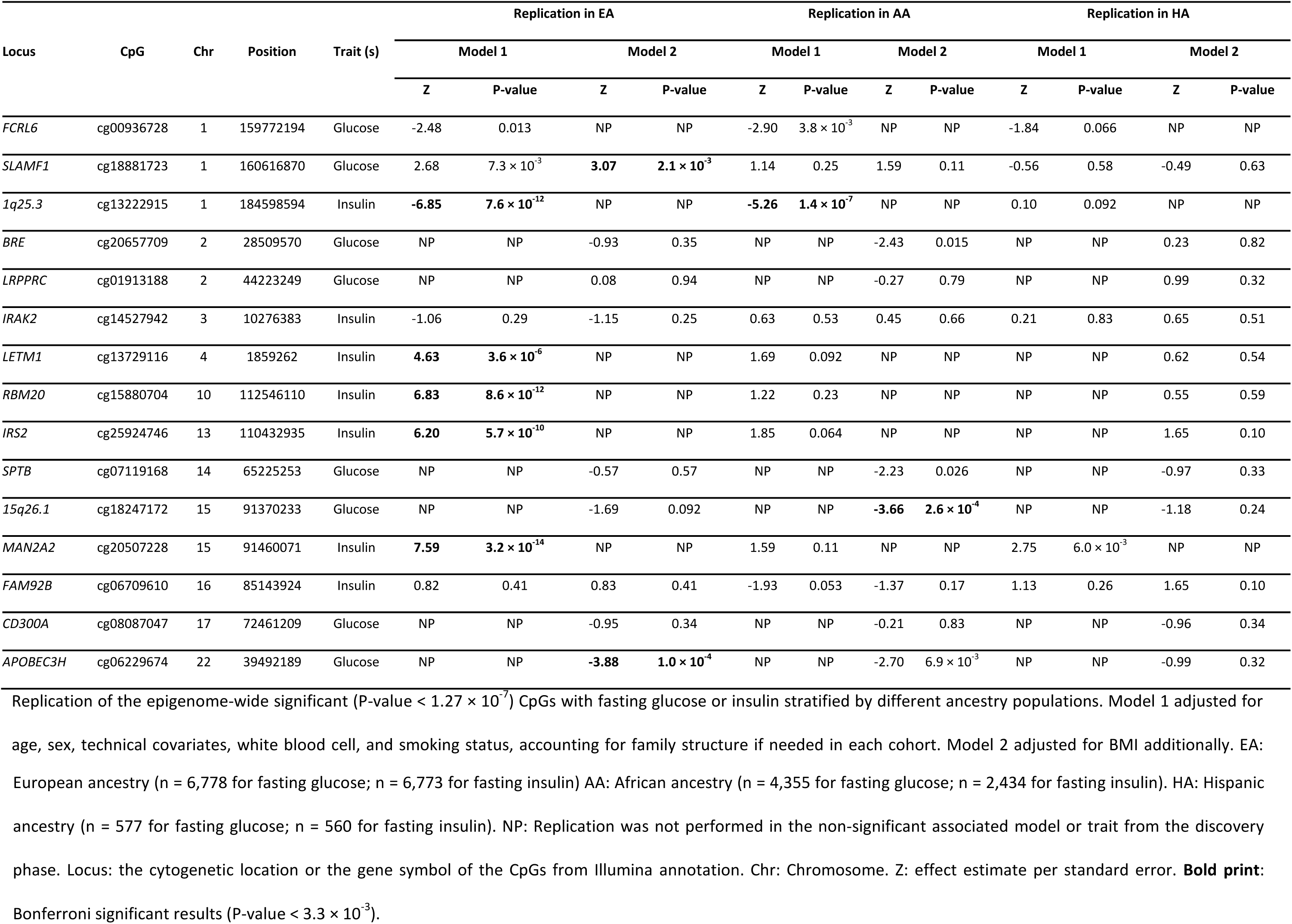
Epigenome-wide association study (EWAS) results: replication of newly discovered differentially methylated sites in different ancestry populations.

**Supplementary Table 4.** Methylation quantitative trait loci (meQTLs) for known or new replicated CpG sites.

| Locus (CpG) | CpG | Variant | Chr | Position | Locus (meQTL) | Type (meQTL) | MAF | Alleles | EA | Z | P-value | Cis/Trans |
| --- | --- | --- | --- | --- | --- | --- | --- | --- | --- | --- | --- | --- |
| *DHCR24* | cg17901584 | rs7412 | 19 | 45412079 | *APOE* | Protein coding | 0.08 | C/T | T | 6.39 | $1.7 \times 10^{-10}$ | Trans |
| *DHCR24* | cg17901584 | rs7701414 | 5 | 131585958 | *P4HA2* | Protein coding | 0.22 | A/G | G | 5.26 | $1.4 \times 10^{-7}$ | Trans |
| *DHCR24* | cg17901584 | rs174550 | 11 | 61571478 | *FADS1/FADS2* | Protein coding | 0.30 | T/C | C | -6.72 | $1.8 \times 10^{-11}$ | Trans |
| *DHCR24* | cg17901584 | rs735665 | 11 | 123361397 | *GRAMD1B*<br>(Nearest) | Protein coding | 0.10 | G/A | A | 10.59 | $3.4 \times 10^{-26}$ | Trans |
| *DHCR24* | cg17901584 | rs687565 | 1 | 55364663 | *TMEM61*<br>(Nearest) | Protein coding | 0.43 | C/A | C | 11.95 | $6.2 \times 10^{-33}$ | Cis |
| *FCRL6* | cg00936728 | rs6657365 | 1 | 159782549 | *FCRL6* | Protein coding | 0.23 | C/G | G | 5.04 | $4.6 \times 10^{-7}$ | Cis |
| *FCRL6* | cg00936728 | rs2523946 | 6 | 29941943 | *HCG9* (Nearest) | lincRNA | 0.49 | C/T | T | 5.33 | $9.8 \times 10^{-8}$ | Trans |
| *SLAMF1* | cg18881723 | rs3129055 | 6 | 29670261 | *ZFP57* (Nearest) | Protein coding | 0.29 | A/G | G | 6.37 | $1.9 \times 10^{-10}$ | Trans |
| *SLAMF1* | cg18881723 | rs11265461 | 1 | 160630143 | *SLAMF1*<br>(Nearest) | Protein coding | 0.36 | C/T | C | 8.39 | $4.7 \times 10^{-17}$ | Cis |
| *1q25.3* | cg13222915 | rs72737737 | 1 | 184598732 | *C1orf21* | Protein coding | 0.06 | A/G | G | 5.68 | $1.4 \times 10^{-8}$ | Cis |
| *4p15.33* | cg10438589 | rs16890352 | 4 | 14385522 | *AC006296.1* | lincRNA | 0.16 | A/G | G | -11.64 | $2.6 \times 10^{-31}$ | Cis |
| *RNF145* | cg26403843 | rs7529925 | 1 | 199007208 | *RP11-16L9.4* | lincRNA | 0.21 | T/C | C | 5.68 | $1.3 \times 10^{-8}$ | Trans |
| *RNF145* | cg26403843 | rs7732603 | 5 | 158614357 | *RNF145* | Protein coding | 0.48 | A/C | C | 35.61 | $8.6 \times 10^{-278}$ | Cis |
| *RBM20* | cg15880704 | rs7906643 | 10 | 112545494 | *RBM20* | Protein coding | 0.09 | C/T | T | -29.45 | $1.4 \times 10^{-190}$ | Cis |
| *CPT1A* | cg00574958 | rs964184 | 11 | 116648917 | *ZNF259* | Protein coding | 0.22 | C/G | G | -5.54 | $3.0 \times 10^{-8}$ | Trans |
| *ASAM* | cg26894079 | rs34817879 | 11 | 123023729 | *CLMP* | Protein coding | 0.12 | T/G | G | -4.67 | $3.0 \times 10^{-6}$ | Cis |
| *KDM2B* | cg13708645 | rs60370741 | 12 | 121966676 | *KDM2B* | Protein coding | 0.30 | T/C | C | 37.16 | $2.6 \times 10^{-302}$ | Cis |
| *IRS2* | cg25924746 | rs9521528 | 13 | 110504805 | *IRS2* (Nearest) | Protein coding | 0.44 | T/A | T | -13.64 | $2.4 \times 10^{-42}$ | Cis |
| *IRS2* | cg25924746 | rs7984800 | 13 | 110671951 | *LINC00396*<br>(Nearest) | lincRNA | 0.26 | A/G | G | 3.88 | $1.0 \times 10^{-4}$ | *Cis* |
| *MYO5C* | cg06192883 | rs1047891 | 2 | 211540507 | *CPS1* | Protein coding | 0.29 | A/C | A | -6.01 | $1.8 \times 10^{-9}$ | *Trans* |
| *MYO5C* | cg06192883 | rs71472932 | 15 | 52541976 | *MYO5C* | Protein coding | 0.11 | G/A | A | 4.22 | $2.4 \times 10^{-5}$ | *Cis* |
| *15q26.1* | cg18247172 | rs404623 | 15 | 91367271 | *BLM* (Nearest) | Protein coding | 0.50 | G/C | G | -11.70 | $1.3 \times 10^{-31}$ | *Cis* |
| *15q26.1* | cg18247172 | rs3129055 | 6 | 29670261 | *ZFP57* (Nearest) | Protein coding | 0.29 | A/G | G | -5.22 | $1.8 \times 10^{-7}$ | *Trans* |
| *15q26.1* | cg18247172 | rs4324798 | 6 | 28776117 | *AL662890.3* | miRNA | 0.04 | G/A | A | -5.48 | $4.3 \times 10^{-8}$ | *Trans* |
| *MAN2A2* | cg20507228 | rs9374080 | 6 | 109616420 | *CCDC162P* | pseudogene | 0.27 | C/T | C | -5.31 | $1.1 \times 10^{-7}$ | *Trans* |
| *MAN2A2* | cg20507228 | rs35831960 | 15 | 91466262 | *MAN2A2*<br>(Nearest) | Protein coding | 0.17 | C/T | T | 8.14 | $4.1 \times 10^{-16}$ | *Cis* |
| *SREBF1* | cg11024682 | rs7701414 | 5 | 131585958 | *P4HA2* | Protein coding | 0.22 | A/G | G | 5.21 | $1.9 \times 10^{-7}$ | *Trans* |
| *SREBF1* | cg11024682 | rs7529925 | 1 | 199007208 | *RP11-16L9.4* | lincRNA | 0.21 | T/C | C | -5.15 | $2.6 \times 10^{-7}$ | *Trans* |
| *SREBF1* | cg11024682 | rs6502629 | 17 | 17869642 | *TOM1L2* | Protein coding | 0.22 | G/A | G | 9.97 | $2.1 \times 10^{-23}$ | *Cis* |
| *TMEM49* | cg24174557 | rs3774937 | 4 | 103434253 | *NFKB1* | Protein coding | 0.25 | C/T | C | -7.75 | $9.3 \times 10^{-15}$ | *Trans* |
| *ABCG1* | cg06500161 | rs225443 | 21 | 43658206 | *ABCG1* | Protein coding | 0.40 | G/A | A | -7.16 | $8.1 \times 10^{-13}$ | *Cis* |
| *APOBEC3H* | cg06229674 | rs28583464 | 22 | 39486593 | *APOBEC3G*<br>(Nearest) | Protein coding | 0.11 | T/C | C | 8.31 | $9.8 \times 10^{-17}$ | *Cis* |

Based on the European blood-based BIOS database (n = 3,841),<sup>26</sup> the meQTL information of known or new replicated CpG sites are shown. Locus (CpG): the cytogenetic location or the gene symbol of the CpGs from Illumina annotation. Locus (meQTL): the located or nearest protein-coding gene of the meQTL from UCSC annotation. Type (meQTL): the gene type of the meQTL. Chr: chromosome. MAF: minor allele frequency. EA: effect allele. Z: effect estimate per standard error.

**Supplementary Table 5.** Common genetic determinants of glycemia related methylation sites and T2D or related traits in blood.

| Variant | Locus<br>(meQTL) | Type (meQTL) | Chr | Position | MAF | EA | Association with CpG |  |  |  | Association with T2D or related traits<sup>†</sup> |  |  |
| --- | --- | --- | --- | --- | --- | --- | --- | --- | --- | --- | --- | --- | --- |
|  |  |  |  |  |  |  | CpG | Locus<br>(CpG) | Z | P-value | Trait | Z | P-value |
| rs6701489 | *TMEM61*<br>(Nearest) | Protein coding | 1 | 55358459 | 0.07 | T | cg17901584 | *DHCR24* | 4.82 | 1.4 × 10<sup>-6</sup> | FG<sup>6</sup> | -3.43 | 8.5 × 10<sup>-4</sup> |
| rs6896438 | *RNF145*<br>(Nearest) | Protein coding | 5 | 158547876 | 0.36 | C | cg26403843 | *RNF145* | 6.15 | 8.0 × 10<sup>-10</sup> | FI<sup>5</sup> | 3.82 | 1.4 × 10<sup>-4</sup> |
| rs10849885 | *KDM2B* | Protein coding | 12 | 121881848 | 0.32 | A | cg13708645 | *KDM2B* | 29.33 | 4.2 × 10<sup>-189</sup> | FG<sup>6</sup> | 4.17 | 2.2 × 10<sup>-5</sup> |
| rs9374080 | *CCDC162P* | pseudogene | 6 | 109616420 | 0.27 | C | cg20507228 | *MAN2A2* | -5.31 | 1.1 × 10<sup>-7</sup> | HbA1c<sup>58</sup> | -5.11 | 2.0 × 10<sup>-7</sup> |
| rs3818717 | *RAI1* | Protein coding | 17 | 17707105 | 0.06 | T | cg11024682 | *SREBF1* | 8.93 | 4.1 × 10<sup>-19</sup> | T2D<sup>27</sup> | 1.08 | 4.9 × 10<sup>-4</sup> |
| rs7529925 | *RP11-16L9.4* | lincRNA | 1 | 199007208 | 0.21 | C | cg11024682 | *SREBF1* | -5.15 | 2.6 × 10<sup>-7</sup> | HbA1c<sup>58</sup> | -3.60 | 2.5 × 10<sup>-4</sup> |
| rs16960744 | *TOM1L2* | Protein coding | 17 | 17755259 | 0.37 | A | cg11024682 | *SREBF1* | 4.87 | 1.1 × 10<sup>-6</sup> | HbA1c<sup>58</sup> | 3.11 | 1.5 × 10<sup>-3</sup> |

The common genetic determinants of glycemia related methylation sites and T2D or related traits are shown. Chr: chromosome. Locus (meQTL): the located or nearest protein-coding gene of the meQTL from UCSC annotation. Type (meQTL): the gene type of the meQTL. MAF: minor allele frequency. EA: effect allele. Locus (CpG): the cytogenetic location or the gene symbol of the CpGs from Illumina annotation. Z: effect estimate per standard error. FG: fasting glucose. FI: fasting insulin. Data sources of associations: 1) association with CpG was from the current discovery phase (n = 4,808), 2) associations with FG (n = 133,010), FI (n = 96,496), T2D (case/control: n = 81,412/370,832) and HbA1c (n = 159,940) were from the MAGIC and DIAGRAM GWAS database<sup>5,6,27,58</sup>

**Supplementary Table 6.**
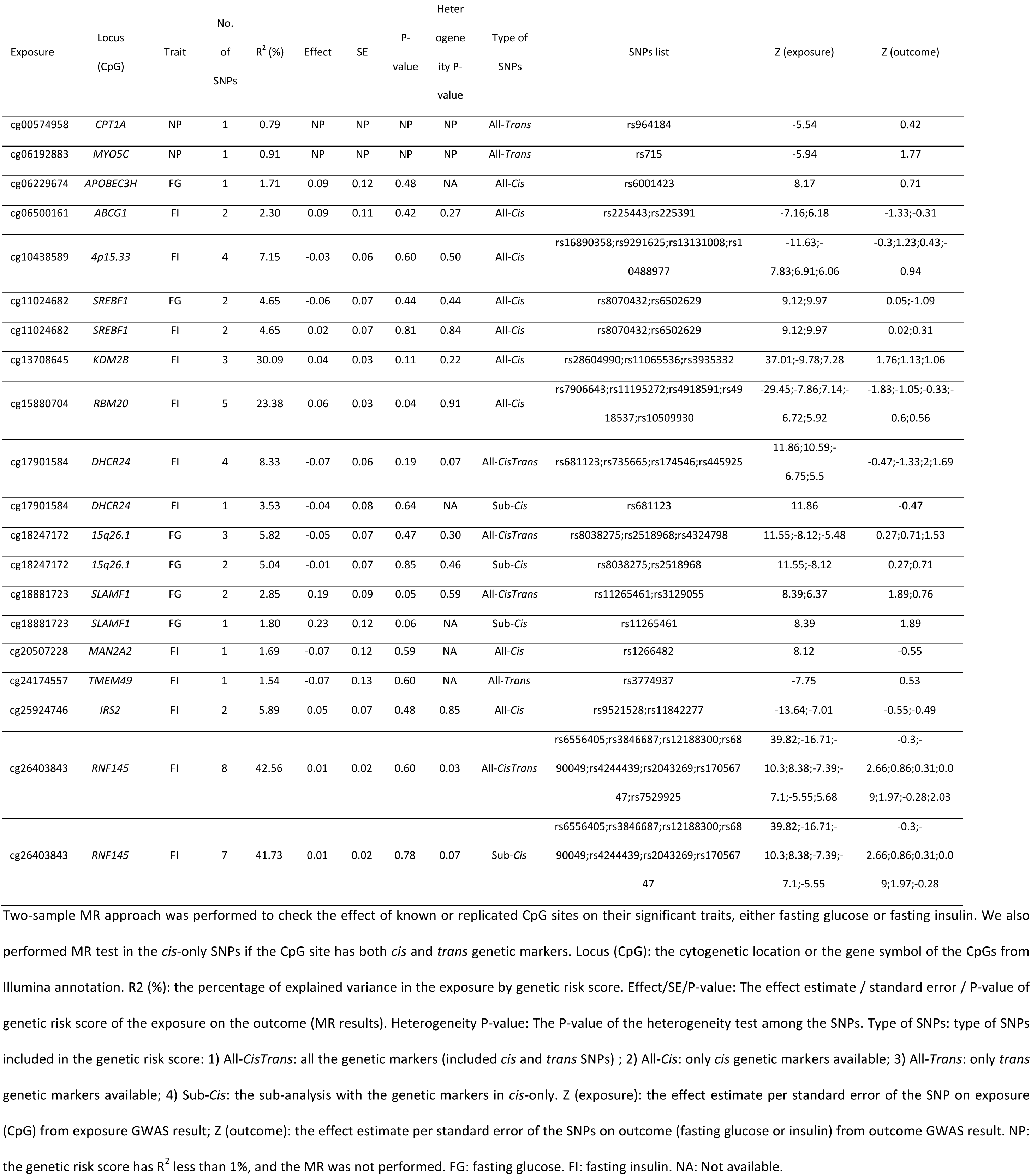
Mendelian randomization (MR) results.

**Supplementary Table 7.** Blood-based expression quantitative trait methylations (eQTMs): association between gene expression and the glycemia related methylation sites.

| Locus (CpG) | CpG | Probe | Probe-Chr | Probe-Pos | Gene expression | Z | P-value | Cis/Trans |
| --- | --- | --- | --- | --- | --- | --- | --- | --- |
| *FCRL6* | cg00936728 | ENSG00000181036 | 1 | 159770301 | *FCRL6* | -13.09 | $4.0 \times 10^{-39}$ | *Cis* |
| *SLAMF1* | cg18881723 | ENSG00000026751 | 1 | 160709037 | *SLAMF7* | 5.84 | $5.4 \times 10^{-9}$ | *Cis* |
| *SLAMF1* | cg18881723 | ENSG00000122223 | 1 | 160832692 | *CD244* | 4.68 | $2.9 \times 10^{-6}$ | *Cis* |
| *SLAMF1* | cg18881723 | ENSG00000117090 | 1 | 160617085 | *SLAMF1* | -4.10 | $4.1 \times 10^{-5}$ | *Cis* |
| *CPT1A* | cg00574958 | ENSG00000110090 | 11 | 68611878 | *CPT1A* | -9.22 | $3.1 \times 10^{-20}$ | *Cis* |
| *SREBF1* | cg11024682 | ENSG00000072310 | 17 | 17740325 | *SREBF1* | -7.84 | $4.5 \times 10^{-15}$ | *Cis* |
| *ABCG1* | cg06500161 | ENSG00000160179 | 21 | 43619799 | *ABCG1* | -12.78 | $2.2 \times 10^{-37}$ | *Cis* |

The gene expressions associated with the glycemia related methylation sites are shown based on the European blood-based BIOS database (n = 3,841) <sup>26</sup>. Locus (CpG): the cytogenetic location or the gene symbol of the CpGs from Illumina annotation. Probe: The probe of the gene expression. Chr: chromosome. Pos: position. Z: effect estimate per standard error.

**Supplementary Table 8.** Common genetic determinants of glycemia related DNA methylation (methylation quantitative trait loci, meQTL) and gene expression (expression quantitative trait loci, eQTL) in blood.

| Variant | Chr | Position | Locus<br>(eQTL) | Type<br>(eQTL) | MAF | EA | Association with CpG |  |  |  | Association with gene expression |  |  |  |  |
| --- | --- | --- | --- | --- | --- | --- | --- | --- | --- | --- | --- | --- | --- | --- | --- |
|  |  |  |  |  |  |  | CpG | Locus<br>(CpG) | Z | P-value | Cis/Trans | Gene<br>expression | Z | P-value | Cis/Trans |
| rs11265282 | 1 | 159774408 | FCRL6 | Protein<br>coding | 0.26 | C | cg00936728 | FCRL6 | 4.17 | $3.0 \times 10^{-5}$ | Cis | FCRL6 | -6.73 | $1.7 \times 10^{-11}$ | Cis |
| rs1577544 | 1 | 160630974 | SLAMF1<br>(Nearest) | Protein<br>coding | 0.39 | T | cg18881723 | SLAMF1 | -5.45 | $5.1 \times 10^{-8}$ | Cis | SLAMF1 | -6.40 | $1.6 \times 10^{-10}$ | Cis |
| rs6502629 | 17 | 17869642 | TOM1L2 | Protein<br>coding | 0.22 | G | cg11024682 | SREBF1 | 9.97 | $2.1 \times 10^{-23}$ | Cis | SREBF1 | -17.93 | $7.2 \times 10^{-72}$ | Cis |

The common genetic determinants of glycemia related methylation sites and gene expression in the European blood-based BIOS database (n = 3,841)<sup>26</sup> are shown. Locus (eQTL): the located or nearest protein-coding gene of the eQTL. Type (eQTL): the gene type of the eQTL. Chr: chromosome. MAF: minor allele frequency. EA: effect allele. Z: effect estimate per standard error. Locus (CpG): the cytogenetic location or the gene symbol of the CpGs from Illumina annotation.

**Supplementary Table 9.**
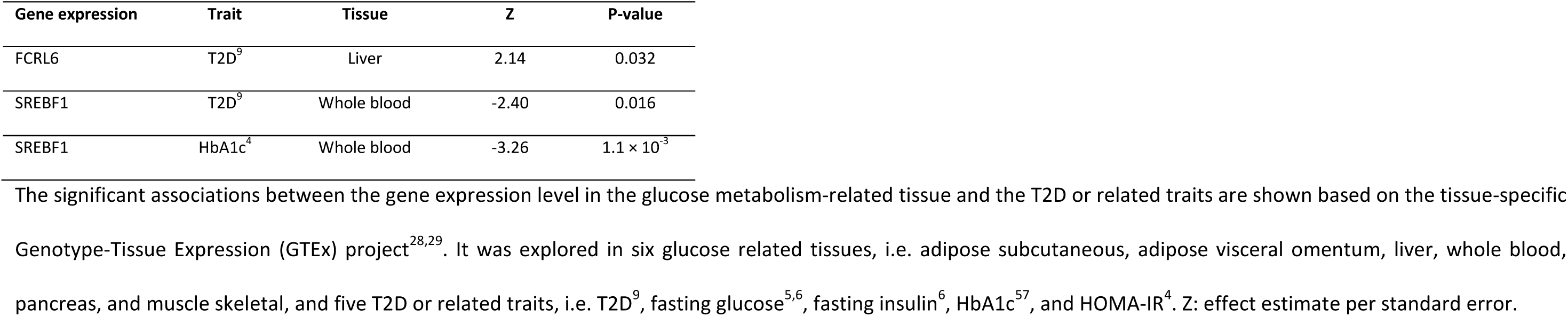
Association between the gene expression level in the glucose metabolism-related tissue and the T2D or related traits based on the Genotype-Tissue Expression (GTEx) project.

| Gene expression | Trait | Tissue | Z | P-value |
| --- | --- | --- | --- | --- |
| FCRL6 | T2D<sup>9</sup> | Liver | 2.14 | 0.032 |
| SREBF1 | T2D<sup>9</sup> | Whole blood | -2.40 | 0.016 |
| SREBF1 | HbA1c<sup>4</sup> | Whole blood | -3.26 | $1.1 \times 10^{-3}$ |

The significant associations between the gene expression level in the glucose metabolism-related tissue and the T2D or related traits are shown based on the tissue-specific Genotype-Tissue Expression (GTEx) project<sup>28,29</sup>. It was explored in six glucose related tissues, i.e. adipose subcutaneous, adipose visceral omentum, liver, whole blood, pancreas, and muscle skeletal, and five T2D or related traits, i.e. T2D<sup>9</sup>, fasting glucose<sup>5,6</sup>, fasting insulin<sup>6</sup>, HbA1c<sup>57</sup>, and HOMA-IR<sup>4</sup>. Z: effect estimate per standard error.

**Supplementary Figure 1.**
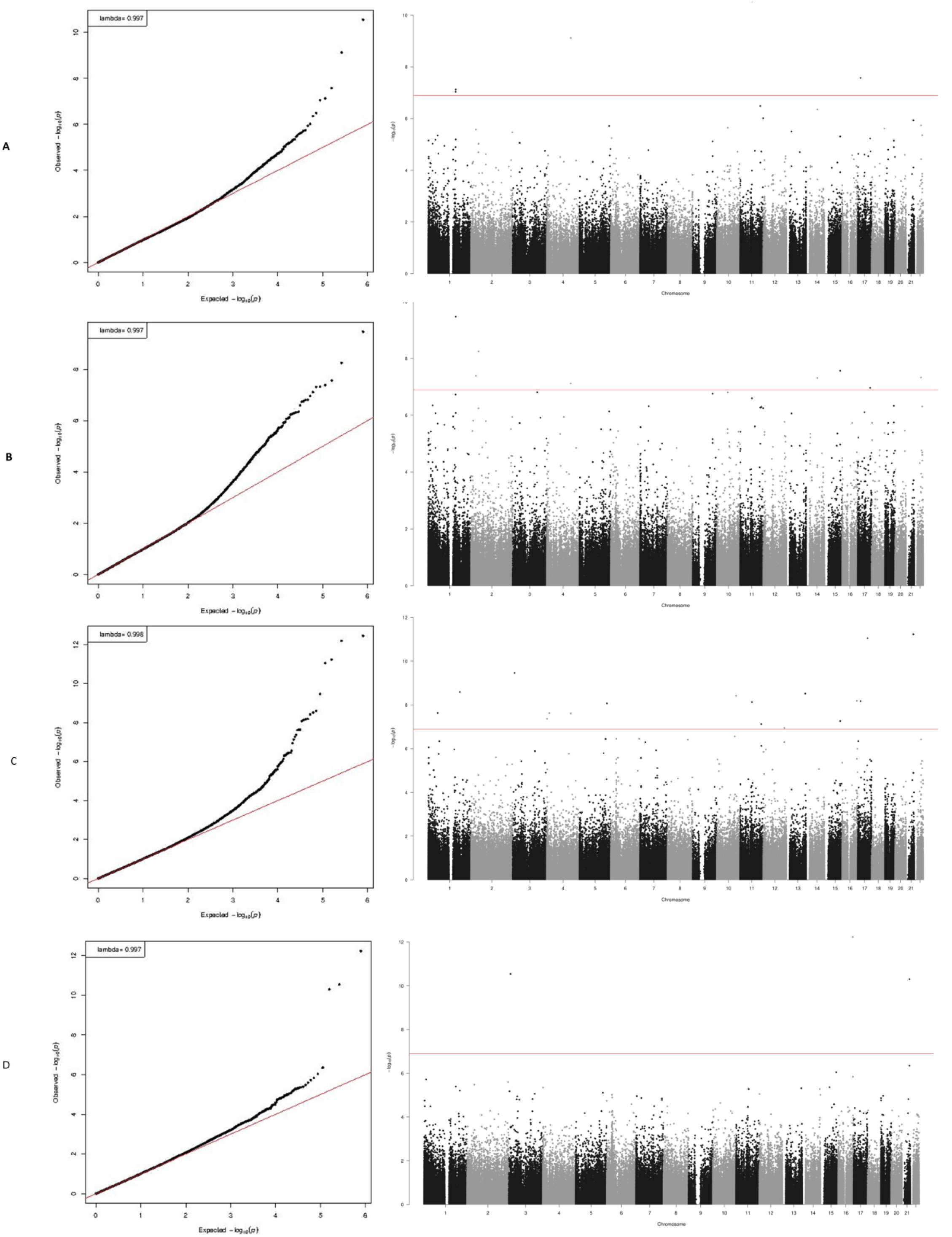
QQ plots and Manhattan plots of the epigenome-wide association study (EWAS) results. A: EWAS results of fasting glucose in the baseline model; B: EWAS results of fasting glucose in the BMI-adjusted model; C: EWAS results of fasting insulin in the baseline model; D: D EWAS results of fasting insulin in the BMI-adjusted model.

**Supplementary Figure 2.**
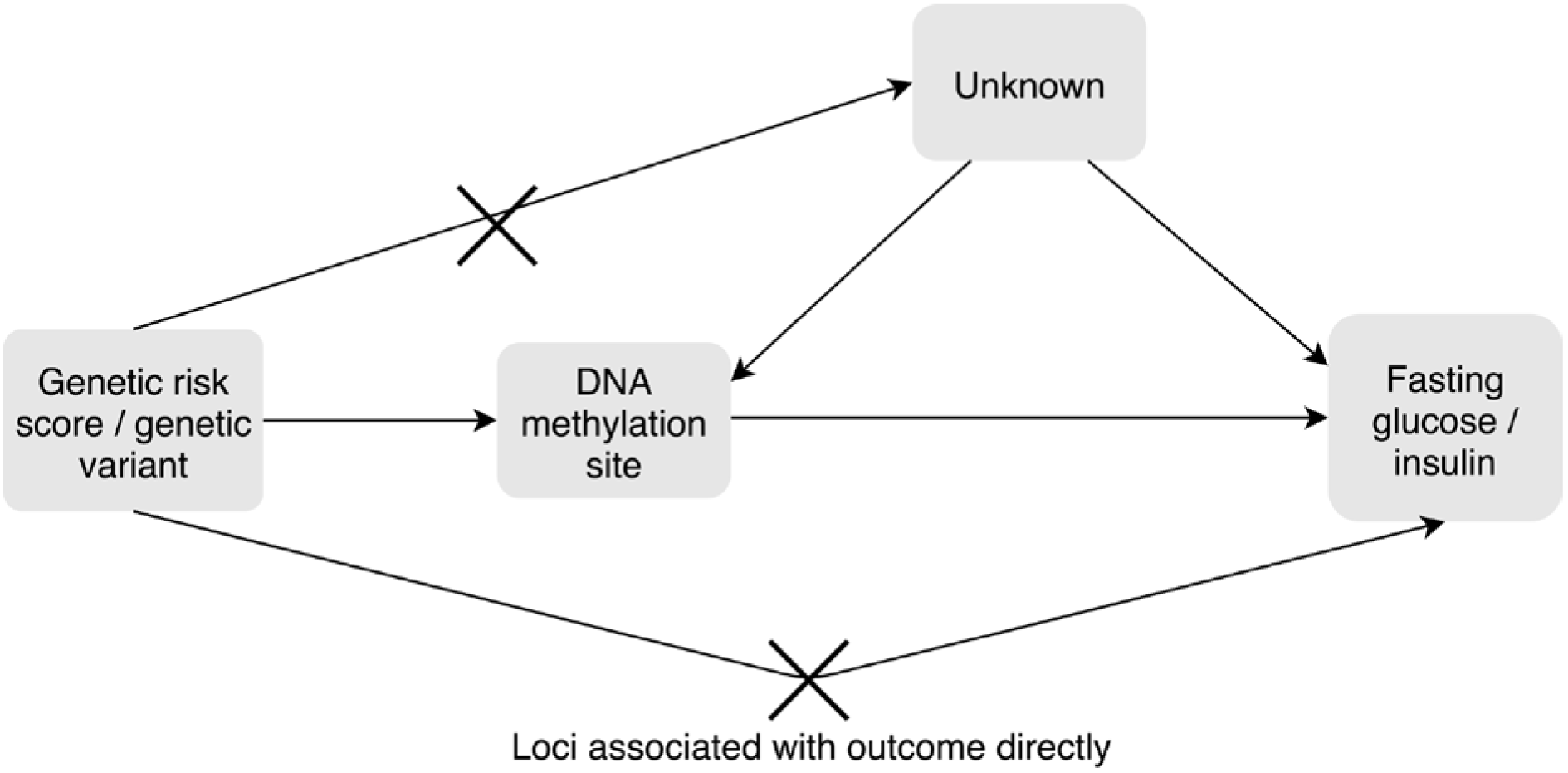
Overview of the general Mendelian Randomization process.

## Disclaimer

The views expressed in this manuscript are those of the authors and do not necessarily represent the views of the National Heart, Lung, and Blood Institute; the National Institutes of Health; or the U.S. Department of Health and Human Services.

